# Long-term exposure to daily ethanol injections in DBA2/J and Swiss mice: Lessons for the interpretation of ethanol sensitization

**DOI:** 10.1101/584102

**Authors:** Vincent Didone, Théo van Ingelgom, Ezio Tirelli, Etienne Quertemont

**Affiliations:** Psychology and Neuroscience of Cognition Research Unit (PsyNCog), *Animal models of cognition*, University of Liège, Liège, Belgium

## Abstract

Most mice ethanol sensitization studies focused on neurobiology at the expense of its behavioral characterization. Furthermore, relatively short ethanol exposures (10 to 20 injections) were used in these studies. The first aim of the present study is to better characterize the development and expression of ethanol sensitization after an extended exposure of 45 daily injections. In some previous studies, mice were classified as “respondent” and “resistant” to ethanol sensitization. The second aim of the present study is to test the long-term reliability of such categorizations and the consequences of their use on the interpretation of the ethanol sensitization results.

Swiss and DBA2/j female mice received 45 consecutive daily ethanol administrations (respectively 2.5 and 2.0 g/kg) and their locomotor activity was daily recorded to test the development of ethanol sensitization. At the end of the procedure, a challenge test assessed the inter-group ethanol sensitization.

The results of the present study show that ethanol sensitization continues to develop beyond 20 days to reach maximal levels after about 25 injections in DBA2/j mice and 40 injections in Swiss mice, although the core phase of the development of ethanol sensitization occurred in both strains during the first 20 days. Remarkably, ethanol sensitization after such a long daily ethanol treatment resulted in both an upward shift of the magnitude of ethanol stimulant effects and a prolongation of these effects in time (up to 30 minutes). Mice classified as “resistant to ethanol sensitization” according to previous studies developed very significant levels of ethanol sensitization when tested after 45 ethanol injections and are best described as showing a delayed development of ethanol sensitization. Furthermore, mice classified as respondent or resistant to ethanol sensitization also differ in their acute response to ethanol, such that it is difficult to ascertain whether these classifications are specifically related to the sensitization process.

## Introduction

Repeated administrations of addictive drugs induce gradual changes in some of their behavioral effects. Some of the drug-induced behaviors decrease over time whereas others gradually increase. The gradual and long lasting behavioral increase is termed “behavioral sensitization”. In laboratory rodents, drug-induced behavioral sensitization is generally modeled as a progressive increase in the locomotor stimulant effects over repeated administrations of the same drug dose (1,2). Chronic exposure to ethanol has been shown to induce a strong and robust sensitization in mice (3,4) and mice are therefore the most widely used animal model to study ethanol sensitization. Ethanol sensitization is believed to result from brain central mechanisms and was suggested to play a key role in alcohol dependence and relapse (2,5,6). Despite some promising results (7,8), such a role of ethanol sensitization clearly needs further confirmations from human studies. Nevertheless, behavioral sensitization is a good experimental tool to explore the neuronal plasticity induced by chronic ethanol exposure.

Many studies were published on the neurobiological bases of ethanol sensitization in mice, involving numerous neurotransmitter systems in the acquisition and expression of ethanol sensitization. However, very few studies were published on the behavioral aspects of ethanol-induced sensitization, especially regarding the procedural and environmental parameters influencing its development and expression. For example, it is unknown how ethanol sensitization evolves when ethanol exposure exceeds three weeks in mice. Indeed, the majority of ethanol sensitization studies used a period of ethanol exposure varying from 10 to 21 days (4,9–21). In the 80’s, some early studies reported extensive periods of ethanol exposure, although they did not use standard sensitization protocol and did not document the daily development of ethanol sensitization. For example, in a study from Masur and Boerngen, no tolerance to ethanol-induced locomotor stimulation was reported after a 60-days ethanol treatment (22). In another study, ethanol was given as the unique source of fluid during 5 months, which was reported to result in an increase of the acute stimulant effects of ethanol (23). Unfortunately, these results were not analyzed and interpreted in terms of behavioral sensitization, because the theories assigning a key role to sensitization in drug addiction were published later (2,6). As a consequence, little is known about the time course of ethanol sensitization after an extensive exposure exceeding a few days. Considering that ethanol consumption in humans usually lasts for years before clear signs of abuse and dependence develop, it seems important to test the effects of longer periods of ethanol administration on the expression of ethanol sensitization in mice. Does ethanol sensitization continue to develop at the same rate after the first few days of ethanol administrations, especially in mice that are categorized as “sensitization resistant”? Does it reach a ceiling effect? Does it even start to decline after reaching a peak level? None of these questions can obtain a response from currently published studies. Therefore, the first aim of the present study was to explore the development of ethanol sensitization over a period of 45 days in which ethanol was daily administered to Swiss and DBA/2j mice, the two most widely used strains of mice in ethanol sensitization studies.

Additionally, a second purpose of the present study was to test whether ethanol sensitization results in a conditioned increase in locomotor activity when sensitized mice are tested after a saline injection in the testing environment. In sensitization studies with psychostimulants, such as cocaine or amphetamines, an excitatory conditioned response is usually observed when mice are tested without drug injection in the testing environment (24,25). Although this explanation remains controversial, such an excitatory conditioned response is often interpreted in Pavlovian terms. The stimulant effects induced by the drug would act as an unconditioned response. After several associations of the test context with drug injections, the environmental cues would become conditioned stimuli that are able to induce an excitatory conditioned response in the absence of drug injection. Although this phenomenon has been largely documented with psychostimulants, the occurrence of an excitatory conditioned response after repeated injections of a stimulant dose of ethanol remains controversial. Whereas a few studies reported excitatory conditioned responses with ethanol, other studies were unable to show such an effect (12,13,26,27). In our laboratory, we never observed an excitatory conditioned response in ethanol sensitization studies (see for example (4)). However, our previous studies tested the excitatory conditioned response after only a few pairings of the test context with ethanol injections (usually 8 to 10 pairings). It remains possible that an excitatory conditioned response with ethanol requires more associations with the test context to develop. Therefore, in the present study, ethanol-sensitized mice from both strains will be tested for an excitatory conditioned response after 46 pairings of the test context with ethanol injections.

Finally, the last aim of the present study was to investigate the reliability of several recent procedures that were used to categorize mice into low and high ethanol sensitized mice or “ethanol sensitization respondent” and “ethanol sensitization resistant” mice. In several published studies, mice were so classified according to the distance travelled during the sensitization sessions after a few days of ethanol injections (10,14,16,19–21,27–40). The categorized groups of mice were then used to study the neurobiological and epigenetic bases of ethanol sensitization vulnerability. For example, Souza-Formigoni and collaborators used an “extreme groups” approach in several studies(14–16,27–29,31–35,37–40). On the basis of their locomotor activity at the 21^th^ ethanol sensitizing session, the upper 33% of mice were classified as “ethanol sensitized” and the lower 33% of mice as “ethanol non-sensitized”. The remaining 34% of mice were not included in the analyses. More recently, the same authors used a median-split technique on the locomotor activity scores at the 21^th^ ethanol sensitization session (30). The lower half of the mice was classified as “low sensitized” whereas the upper half was classified as “high sensitized”. Finally, Botia and collaborators classified their mice as “sensitized” or “resistant” based on a ratio between the 10^th^ and the first session of sensitization (10,19–21) using a more complex formula (see below). In the present study, the statistical implications of such categorizations of continuous variables will not be discussed, although it is worth mentioning that statisticians usually do not recommend splitting continuous variables into categories and instead advise the use of proper statistical methods for metric variables, such as correlations or multiple regression analyses (41–44). The purpose of the present study is to explore the reliability of such categorization and the consequences of their use on the interpretation of the results. In particular, we were interested in studying whether mice classified as “resistant” or “non-sensitized” on the basis of their levels of sensitization on the 10^th^ or the 21^th^ ethanol session respectively remain non-sensitized after 45 ethanol injections. Indeed, it is possible that the so called “sensitization resistant” mice simply develop ethanol sensitization later. They might even develop higher levels of ethanol sensitization after 45 ethanol injections. Conversely, the sensitized mice might reach a ceiling level of sensitization or even show a subsequent decrease in ethanol sensitization expression. Such a pattern of results would strongly affect the interpretation of the previously published results. For example, mice previously classified as “sensitized” might be reclassified as “rapidly sensitized” or even “temporarily sensitized”, whereas “non-sensitized” mice might be characterized as “delayed sensitization” or “postponed sensitization” according to the present results.

## Material and methods

### Animals

For the whole study, 95 Swiss and 65 DBA/2J female mice were used. Swiss mice were bred in our colony from progenitors purchased from Janvier Laboratories (Le Genet St Isle, France). Female DBA/2J mice were purchased from Janvier Laboratories. Animals were 10 weeks old at the beginning of the experiments. One week before the tests, mice were housed two per cage in transparent polycarbonate cages (332 × 150 × 130mm) provided with pine sawdust bedding. They had access to water and food (standard pellets, Carfil Quality, BVDA, Oud-Turnhout, Belgium) ad libitum except during the experimental procedures. The colony room was kept on a 12:12h light-dark cycle (light on at 6 a.m.). The experimental and the colony rooms were maintained on an ambient temperature of 22±1°C. All experimental procedures were carried out between 7 a.m. and 2 p.m. All experimental treatments and animal maintenance were reviewed by the University of Liege Animal Care and Experimentation Committee, which gave its approval according to the Belgian implementation of the animal welfare guidelines laid down by the European Union (“Directive 2010/63/EU of the European Parliament and of the Council of 22 September 2010 on the protection of animals used for scientific purposes”).

### Drugs

Absolute ethanol (99.99%) was purchased from VWR International (Leuven, Belgium) and diluted (20% v/v) in an isotonic 0.9% saline solution. All injections were administered via the intraperitoneal (i.p.) route.

### Behavioral sensitization

The same experimental protocol was used in all experiments to induce and test the behavioral sensitization to ethanol in the two strains of mice. The experimental protocol was adapted from previous ethanol sensitization studies from our laboratory (3,4,9). On the first day of the experiment (P0, acclimation session), mice were acclimated to the apparatus and experimental procedure. All mice were moved to the experimental room, weighed, injected with saline, and immediately placed into the open-fields (40×40×40 cm). The open-fields have black Plexiglas walls and floor. The distance travelled (cm) was recorded for the next 5 min by a computer using an activity tracking system, Videotracking® (France, Lyon). The sensitization procedure started the next day. During 45 consecutive days (days 1-45), mice were moved to the experimental room, weighed, injected with ethanol (or saline for the control groups), and immediately placed into the experimental chambers. Swiss mice were injected with 2.5 g/kg ethanol whereas DBA2/J mice were injected with 2.0 g/kg ethanol. These doses were selected as the optimal doses for inducing ethanol sensitization in these two strains of mice on the basis of previously published studies (4,9,10,18,20,21). Pilot studies from our laboratory also confirmed slight differences between strains of mice in the proper ethanol doses to obtain a significant behavioral sensitization (unpublished data). After ethanol injection, the distance travelled (cm) by mice was then recorded for 5 min. This session length was chosen in order to specifically capture the stimulant effects occurring during the ascending limb of the blood alcohol concentration. The expression of ethanol sensitization was assessed on the 46th day. All mice were injected with ethanol (respectively 2.5 and 2.0 g/kg for Swiss and DBA2/J mice) and immediately placed into the experimental chambers. Their locomotor activity was recorded for 30 min. The longer duration of this session was used to capture a possible delayed effect. On the next day, all the mice were tested for their conditioned locomotor response after a saline challenge (47^th^ day) in the testing environment. All the mice were injected with saline and immediately placed into the experimental chambers. Their locomotor activity was recorded for 30 min.

### Classification of chronically ethanol-treated mice

DBA2/J and Swiss mice from ethanol groups were classified as “resistant” or as “respondent” to ethanol sensitization according to the criteria used in previously published studies (10,14,16,19–21,27–40). The first classification was a median split of the mice according to their locomotor response on the 21^st^ ethanol session following the procedure from Ferreira and colleagues (30). The upper 50% of the mice with the highest ethanol-induced locomotor activity were defined as “respondent to ethanol sensitization”, whereas the lower 50% of the mice were defined as “resistant to ethanol sensitization”. The second classification was the extreme group procedure used by Souza-Formigoni and colleagues (14–16,27–29,31–35,37–40). Mice were classified according to their locomotor activity on the 21^st^ ethanol session. The upper 33% of the mice with the highest ethanol-induced locomotor activity were defined as “respondent to ethanol sensitization”, whereas the lower 33% of the mice were defined as “resistant to ethanol sensitization”. The remaining 34% of the mice were not included in the analysis. Finally, the third classification followed the procedure used by Botia and colleagues (10,19–21). A within-subject sensitization score was calculated for each mouse as the ratio between locomotor activity on the 10^th^ ethanol session and on the 1^st^ ethanol session (Day10/Day1 ratio). Mice were defined as “respondent to ethanol sensitization” if the percentage of increase in that ratio exceeded the coefficient of variance (CV) of the control group. According to the authors, this CV provided a measure of locomotor variability over the course of the experiment undue to ethanol administrations. For example, a Swiss mouse with a 1.26 sensitization ratio shows a 26% increase in locomotor activity and would meet the criteria if the CV of the saline control group is 0.25 or lower. Mice that did not fulfill this criterion were classified as “resistant to ethanol sensitization”.

### Statistics

Analyses of variance (ANOVAs) for mixed designs were computed on the travelled distances during the sensitization sessions (days 1-45) to test for the development of ethanol sensitization in each strain of mice separately. The sensitization sessions were defined as a within-subject factor, while the treatment (ethanol vs. saline) was defined as a between-subject factor. Additionally, planned contrasts were computed independently for each group of mice to compare their locomotor activity on the 1^st^, the 10^th^, the 21^st^ and 45^th^ sessions in order to identify significant within-subject ethanol sensitization or habituation to saline administrations. The 10^th^ and 21^th^ sessions were selected as they were the last ethanol sessions in the classification studies according to which mice were classified as “respondent to ethanol sensitization” or “resistant to ethanol sensitization” (10,30,40).

The results of the ethanol (46^th^ day) and saline (47^th^ day) test sessions were analyzed with mixed design ANOVAs in which the treatment administered during the acquisition phase was defined as a between-subject factor (ethanol vs. saline) and the 5 min time interval as within-subject factor (6 levels). Effect sizes were computed as simple eta-squared (η²) for each tested effect.

Tukey’s HSD post-hoc tests were performed to further investigate mean differences between groups.

Mixed design ANOVAs were computed to test for differences in the development of ethanol sensitization between mice from both strains that were classified as “respondent to ethanol sensitization” or “resistant to ethanol sensitization” according to the three classification procedures described above. The sensitization sessions were defined as a within-subject factor, while the group (resistant, respondent and control) was defined as a between-subject factor. To further investigate group differences, planned contrasts were computed to compare the locomotor activity of the three groups on the 1^st^ and 45^th^ sensitization sessions. Additionally, planned contrasts were computed independently for each group of mice to compare their locomotor activity on the 1^st^ and 45^th^ sensitization sessions in order to identify significant within-subject ethanol sensitization or habituation to saline administrations. Similar number of resistant and respondent mice would be obtained whatever the moment of the classification for the first two classifications (50% of the mice for the median split and 33% of the mice for the extreme group procedure). However, the number of respondent mice is likely to evolve with time when mice are classified using the third classification procedure from Botia and colleagues (10,19–21). Therefore, in order to test for such an evolution, mice were reclassified using this same procedure according to their ethanol-induced locomotor activity on the 10^th^, 21^st^ and 45^th^ ethanol sessions. A Cochran’s Q test was then used to test whether the percentage of respondent/resistant mice changed over ethanol sessions using this categorization procedure.

For all the experiments, the assumption of homogeneity of variances was assessed using a Levene’s test. When required, square root transformations were used to normalize the data before the ANOVAs. However, for the sake of clarity, means of the raw values are presented in the figures. Statistical significance was set at P<0.05.

## Results

### Behavioral sensitization

#### Development and expression of sensitization in female Swiss mice

As shown in Fig. 1, the locomotor activity of female Swiss mice gradually increases over repeated administrations of ethanol, whereas the saline control group remains stable. This is confirmed by significant main effects for the group (ethanol vs. saline; F_1,93_=87.13, p<0.0001, η²=0.41), and for the session (F_44,4092_=23.99, p<0.0001, η²=0.063), together with a significant interaction between the two variables (F_44,4092_=30.17, p<0.0001, η²=0.080). Several stages in the development of a locomotor sensitization to ethanol can be visualized on Fig. 1. There was a sustained and constant increase in locomotor activity during the first 21 days, followed by a slower increase to reach the highest levels of activity between days 35 and 45. The planned contrasts show a significant difference between the acute response to ethanol (day 1) and the locomotor response to ethanol at the 10^th^ (p<0.0001) and 21^st^ (p<0.0001) ethanol sessions. On the last ethanol session (day 45), locomotor activity was significantly higher (p<0.0001) than on the 1^st^, 10^th^ and 21^st^ ethanol sessions. In contrast, locomotor activity did not significantly differ on those days in the saline control group.

**Fig 1.**
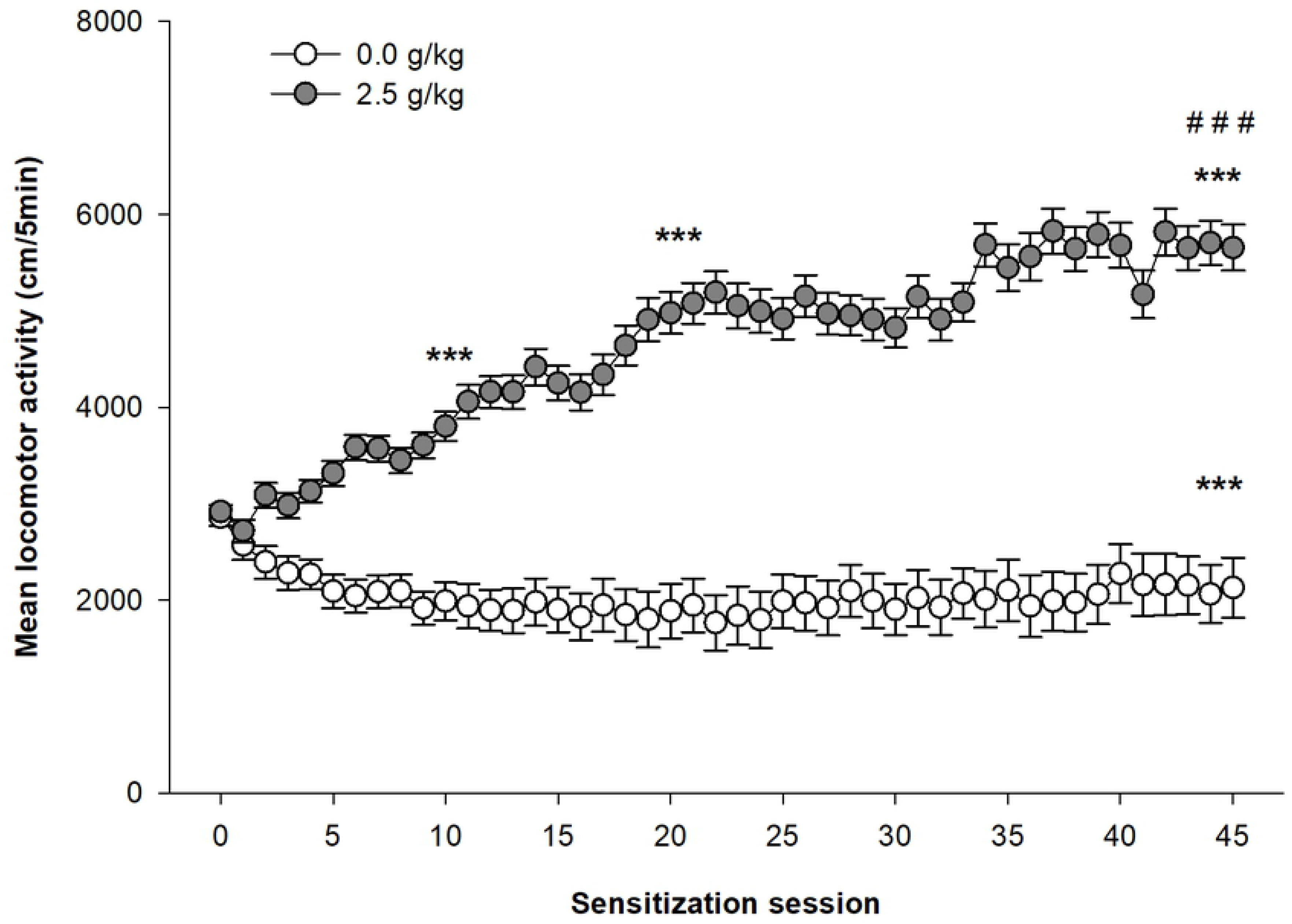
Development of sensitization to ethanol in female Swiss mice. Female Swiss mice were injected daily with ethanol (2.5g/kg) or saline. Data are expressed as mean ± SEM. ***p<0.0001: significantly different from the first acquisition session, as indicated by the planned contrasts. ###p<0.0001: significantly different from all the others sessions as indicated by the planned contrasts.

Figures 2A and 2B respectively show the time course of locomotor activity and the mean distance travelled during the six 5-min intervals of the test session after 45 daily ethanol administrations. All mice were injected with 2.5g/kg ethanol before the test session. The mixed design ANOVA computed on the distances travelled by the two groups of mice on the test session shows significant main effects for the group (ethanol vs. saline; F_1,93_=135.31, p<0.0001, η²=0,51) and for the time intervals (F_5,465_=14.48, p<0.0001, η²=0,044). The interaction group x time intervals was also significant (F_5,465_=2,24, p=0.049, η²=0.0071). The Tukey’s HSD tests confirm that ethanol-sensitized mice show significantly higher levels of locomotor activity than the control group over all six-time intervals of the test session. The significant interaction group x time intervals is also clarified by the post-hoc tests. In ethanol-sensitized mice, the stimulant effects of ethanol were stable during the 30-min duration of the test session, as confirmed by the lack of significant differences between the six time intervals on the post-hoc tests. In contrast, ethanol-induced locomotor activity in the control group adopts the usual pattern of acute stimulant ethanol effects in Swiss mice: a brief significant increase in locomotor activity during 5-10 minutes followed by a return to basal levels of activity. This effect is confirmed by the post-hoc tests showing a significant increase in locomotor activity in that group during the first two 5-min intervals (see Fig. 2A).

**Fig. 2.**
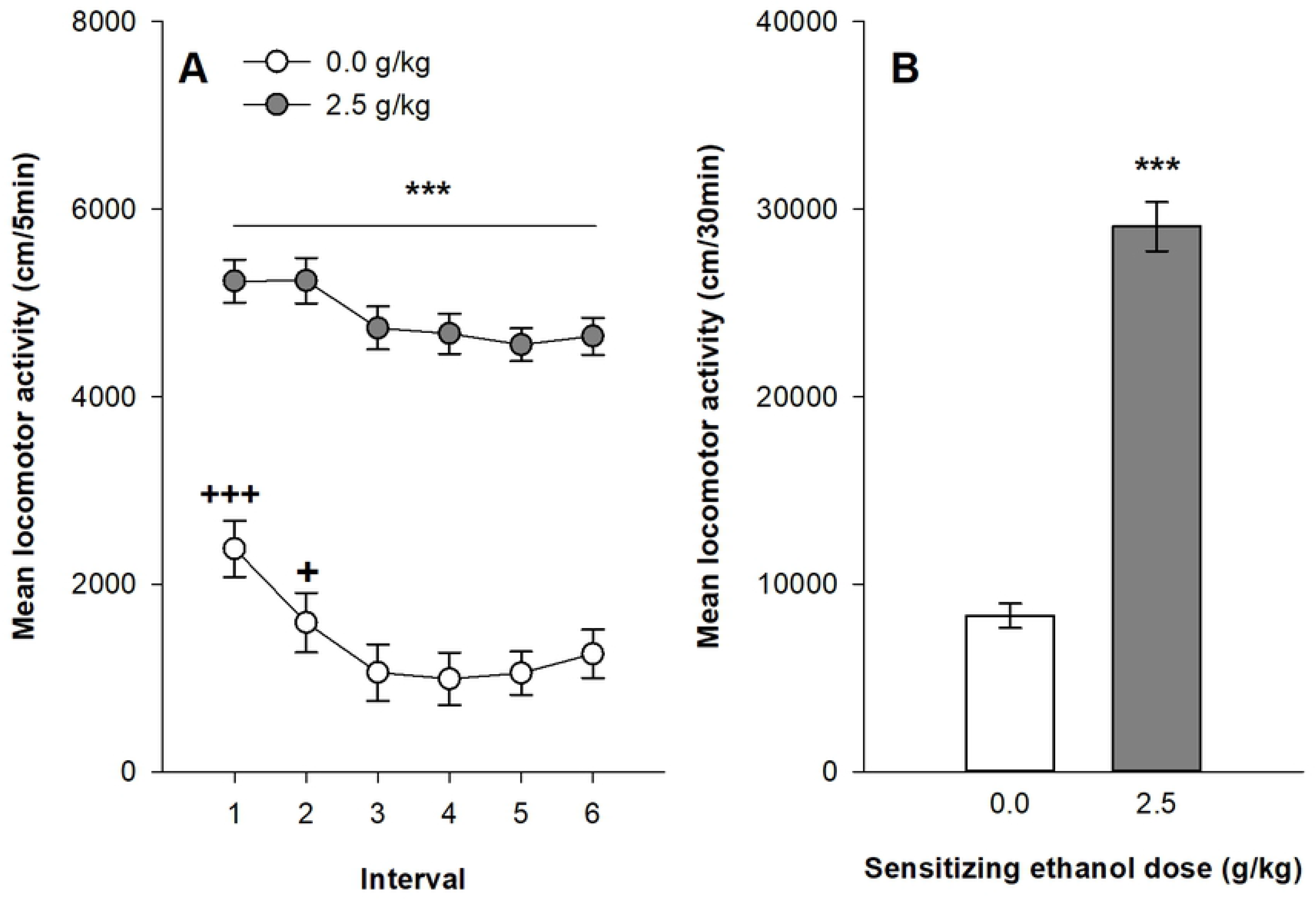
Expression of behavioral sensitization following 45 ethanol administrations (2.5 g/kg) in Swiss female mice. Data are expressed as mean ± SEM. (A) depicts the time course for expression of sensitization following a 2.5 g/kg ethanol injection in mice that were sensitized with ethanol or saline. (B) shows the total distance travelled during the 30 min test by mice sensitized or not to ethanol. +p<0.05; +++p<0.0001: significantly different of the other session within the same experimental condition. ***p<0.0001: significantly different from the respective control group that was repeatedly injected with saline during the acquisition sessions, as indicated by the Tukey HSD post hoc test.

#### Conditioned response in female Swiss mice

On the day following the ethanol sensitization test session, all mice were tested for their conditioned locomotor response after a saline challenge. The mixed design ANOVA computed on the distances travelled during the six 5-min intervals of the session shows significant main effects for the group (ethanol vs. saline; F_1,93_=15.76, p<0.0001, η²=0.12) and for the time intervals (F_5,465_=191.55, p<0.0001, η²=0.32). There is also a significant interaction group x time intervals (F_5,465_=33.74, p<0.0001, η²=0.077). As shown on figure 3, ethanol-sensitized mice express higher levels of locomotor activity than mice from the control group. Post–hoc analyses indicate that the differences between the groups are limited to the first ten minutes of the session. The distances travelled during the last 20 minutes of the session were virtually equal in both groups. As these results might indicate either an excitatory conditioned response or a lack of habituation to the test context in ethanol-sensitized mice, further analyses were performed to test those hypotheses. A mixed design ANOVA was computed on the locomotor activity of both groups during both the habituation session (day 0) and conditioned test session (day 47). This ANOVA shows a significant interaction group x session (F_1,93_=32.95, p<0.0001, η²=0.12). The post-hoc tests indicate that locomotor activity between the habituation and conditioned test sessions decreased in the control group, whereas it remained virtually unchanged in ethanol-sensitized mice, supporting the lack of habituation hypothesis.

**Fig 3.**
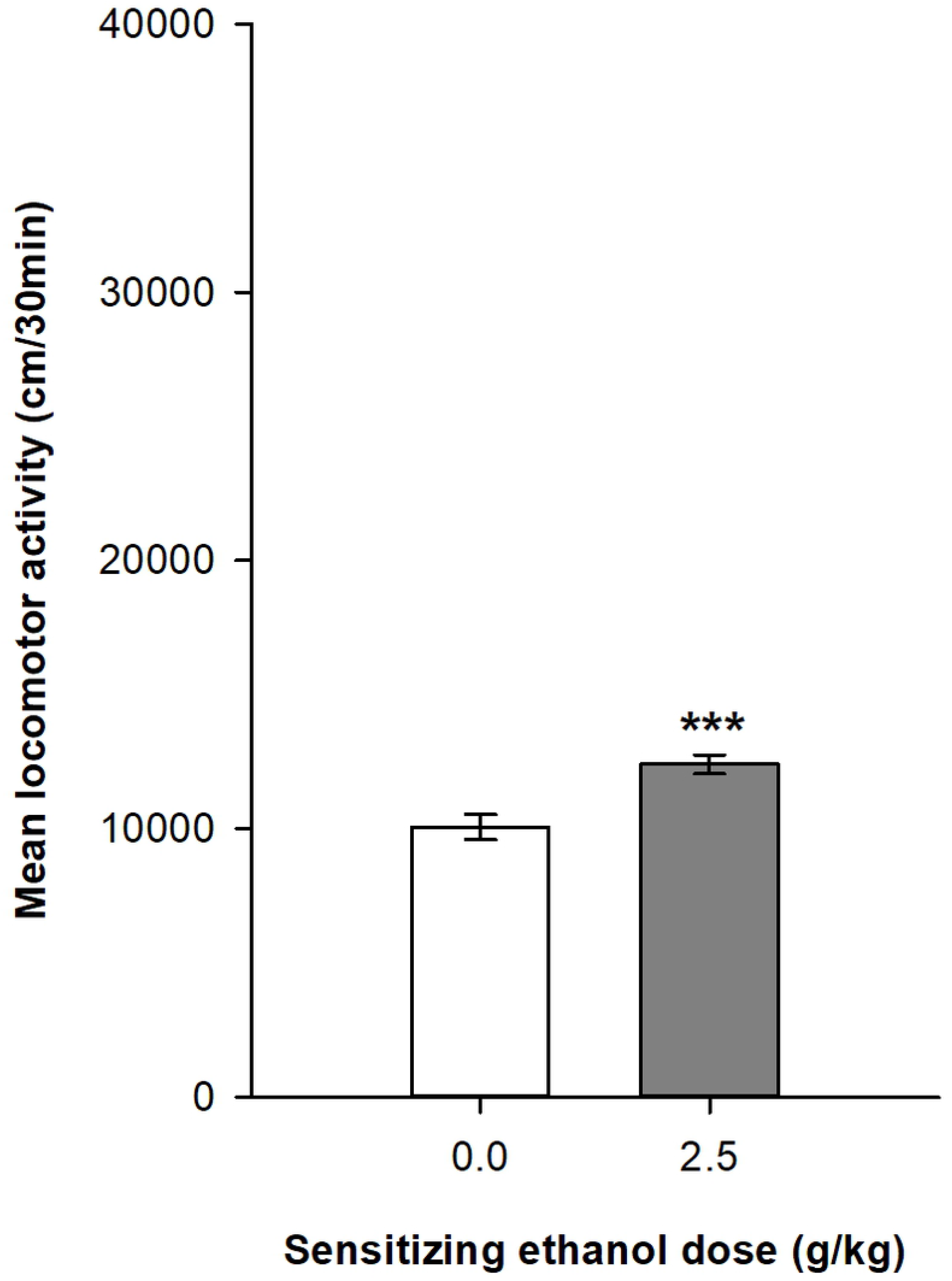
Total distance travelled during the conditioned response test in female Swiss mice. Female Swiss mice were administered with saline to evaluate the conditioned response induced by chronic ethanol administration (2.5 g/kg). ***p<0.0001: significantly different from the control group that was repeatedly injected with saline during the acquisition sessions.

#### Development and expression of sensitization in female DBA2/j mice

The mixed design ANOVA computed on the distances travelled in the 45 repeated ethanol sessions shows significant main effects for the group (ethanol vs. saline; F_1,63_=64.13, p<0.0001, η²=0.38) and for the session (F_44,2772_=6.43, p<0.0001, η²=0.040). There is also a significant interaction group x session (F_44,2772_=3.31, p<0.00001, η²=0.021) which is the result of the gradual increase of locomotor activity over the sessions in the ethanol group, whereas the saline control group remains relatively stable (see Figure 4). The locomotor activity progressively increases with ethanol sessions to reach a maximum at the 21^st^ session. The planned contrasts computed to test for differences between the 1^st^, 10^th^, 21^st^ and 45^th^ sessions indicate significant differences between the 1^st^ acute sessions and all other sessions. There are also significant differences between the 10^th^ session and the 21^st^ and 45^th^ sessions, whereas these latter two sessions do not significantly differ. None of those differences are statistically significant in the saline control group.

**Fig 4.**
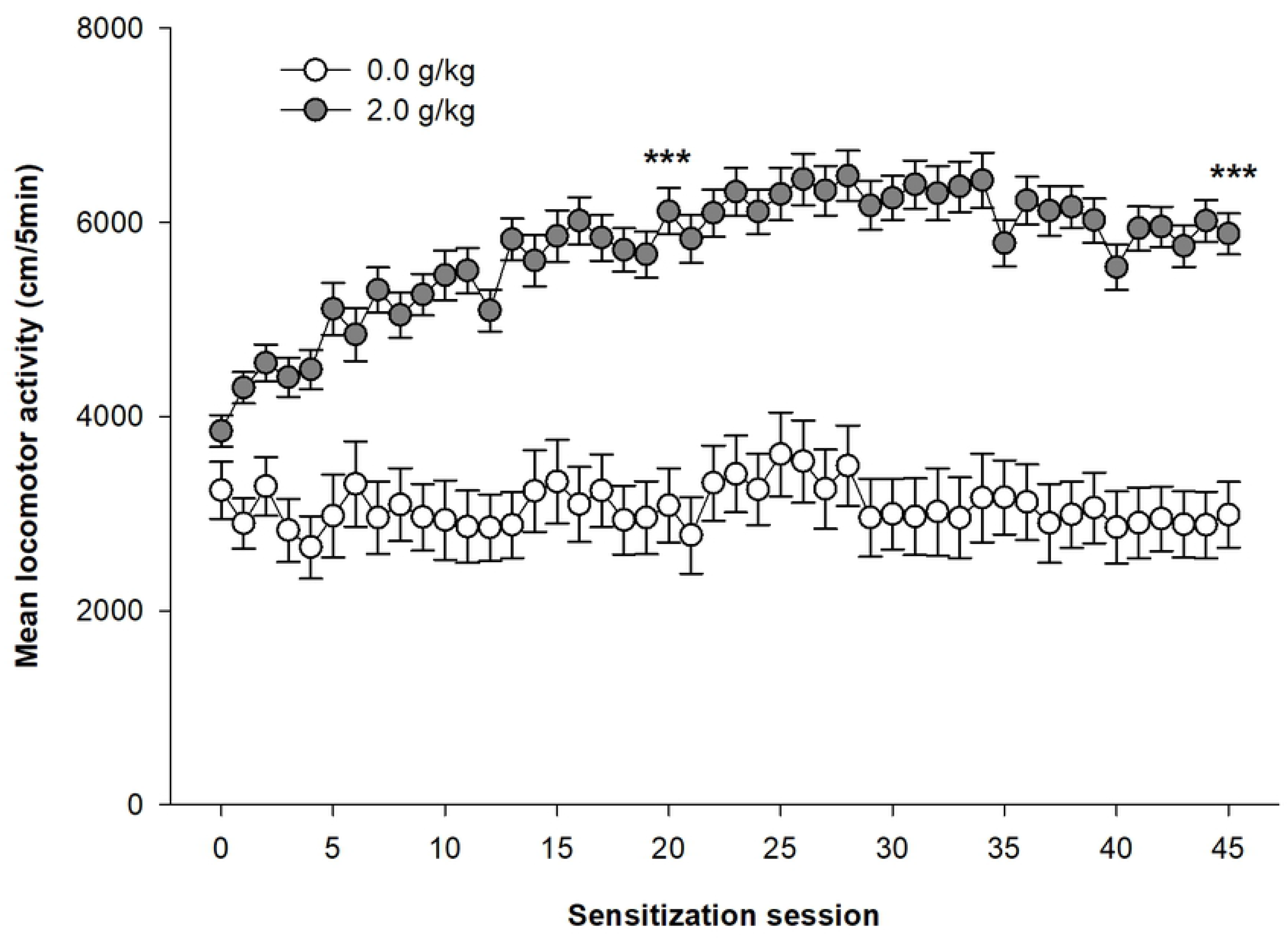
Development of sensitization to ethanol in Female DBA/2j mice. Female DBA/2j mice were injected daily with ethanol (2.0 g/kg) or saline. Data are expressed as mean ± SEM. ***p<0.0001: significantly different from the first acquisition session, as indicated by the planned contrasts.

Figure 5 show the expression of behavioral sensitization in female DBA2/j mice after 45 daily ethanol administrations. All mice were injected with 2.0 g/kg ethanol before the test session. The mixed design ANOVA computed on the distances travelled by the two groups of mice on the test session shows significant main effects for the group (ethanol vs. saline; F_1,63_=23.16, p<0.0001, η²=0.17) and for the time intervals (F_5,315_=4.58, p<0.001, η²=0.031). In contrast to the results in Swiss mice, the interaction group x time intervals was not significant (F_5,315_=1.03, p=0.397, η²=0.007). The Tukey’s HSD tests confirm that ethanol-sensitized mice show significantly higher levels of locomotor activity than the control group over all six time intervals of the test session.

**Fig 5.**
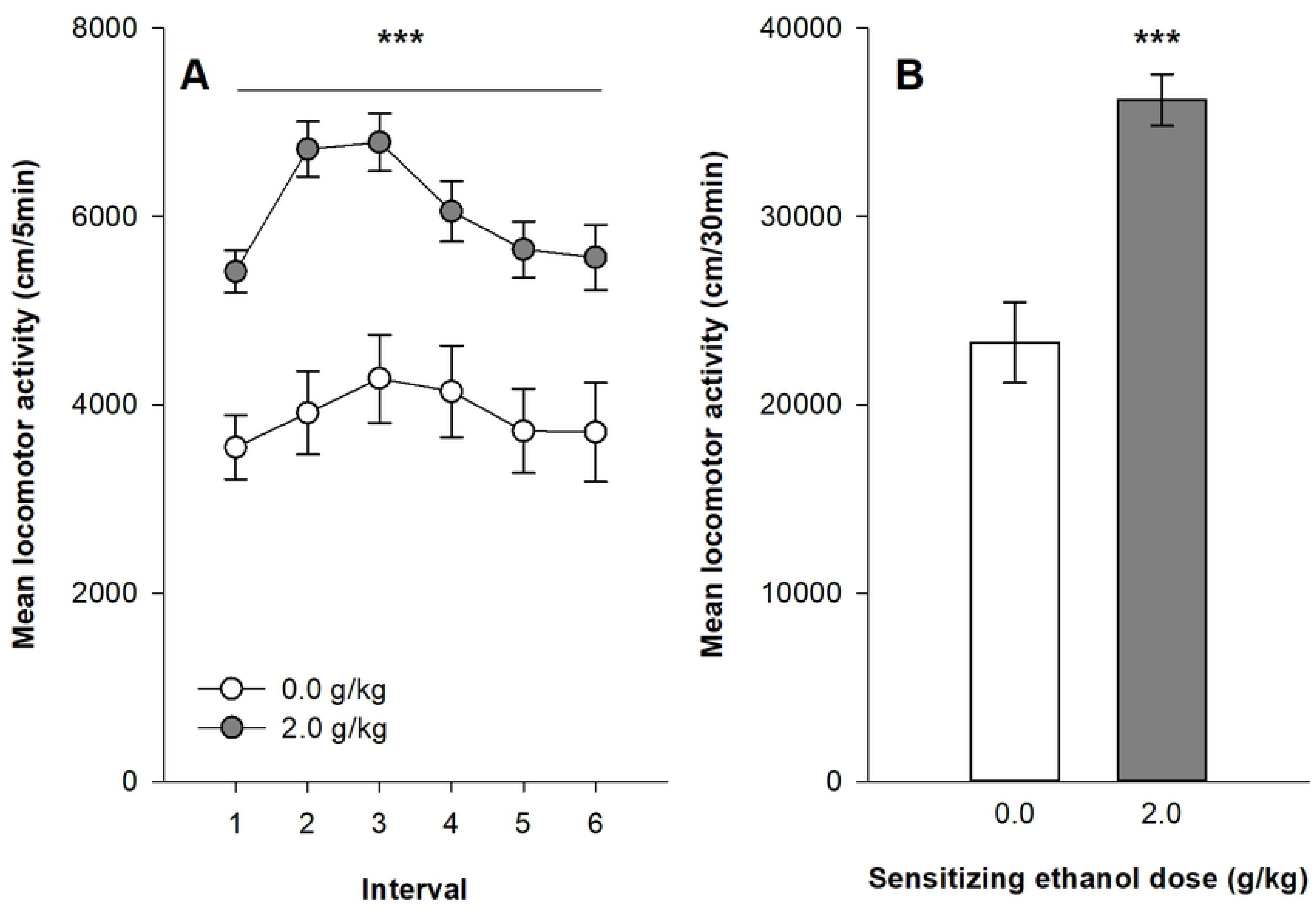
Expression of behavioral sensitization following 45 ethanol administrations (2.0 g/kg) in DBA2/j female mice. Data are expressed as mean ± SEM. (A) depicts the time course for expression of sensitization following a 2.0 g/kg ethanol injection in mice that were sensitized with ethanol or saline. (B) shows the total distance travelled during the 30 min test of expression by mice sensitized or not to ethanol. ***p<0.0001: significantly different from the respective control group that was repeatedly injected with saline during the acquisition sessions, as indicated by the Tukey HSD post hoc test.

#### Conditioned response in female DBA2/j mice

On the day following the ethanol sensitization test session, all mice were tested for their conditioned locomotor response after a saline challenge. The mixed design ANOVA computed on the distances travelled during the six 5-min intervals of the session shows no significant main effects for the group (ethanol vs. saline; F_1,62_=0.78, p=0.38, η²=0.0099) or the time intervals (F_5,310_=1.61, p=0.158, η²=0.0064). Additionally, there is no significant interaction group x time intervals (F_5,310_=1.61, p=0.158, η²=0.0052). As shown on Figure 6, there are no differences in mean locomotor activity between the two groups, which indicates the lack of conditioned response in DBA2/J mice.

**Fig 6.**
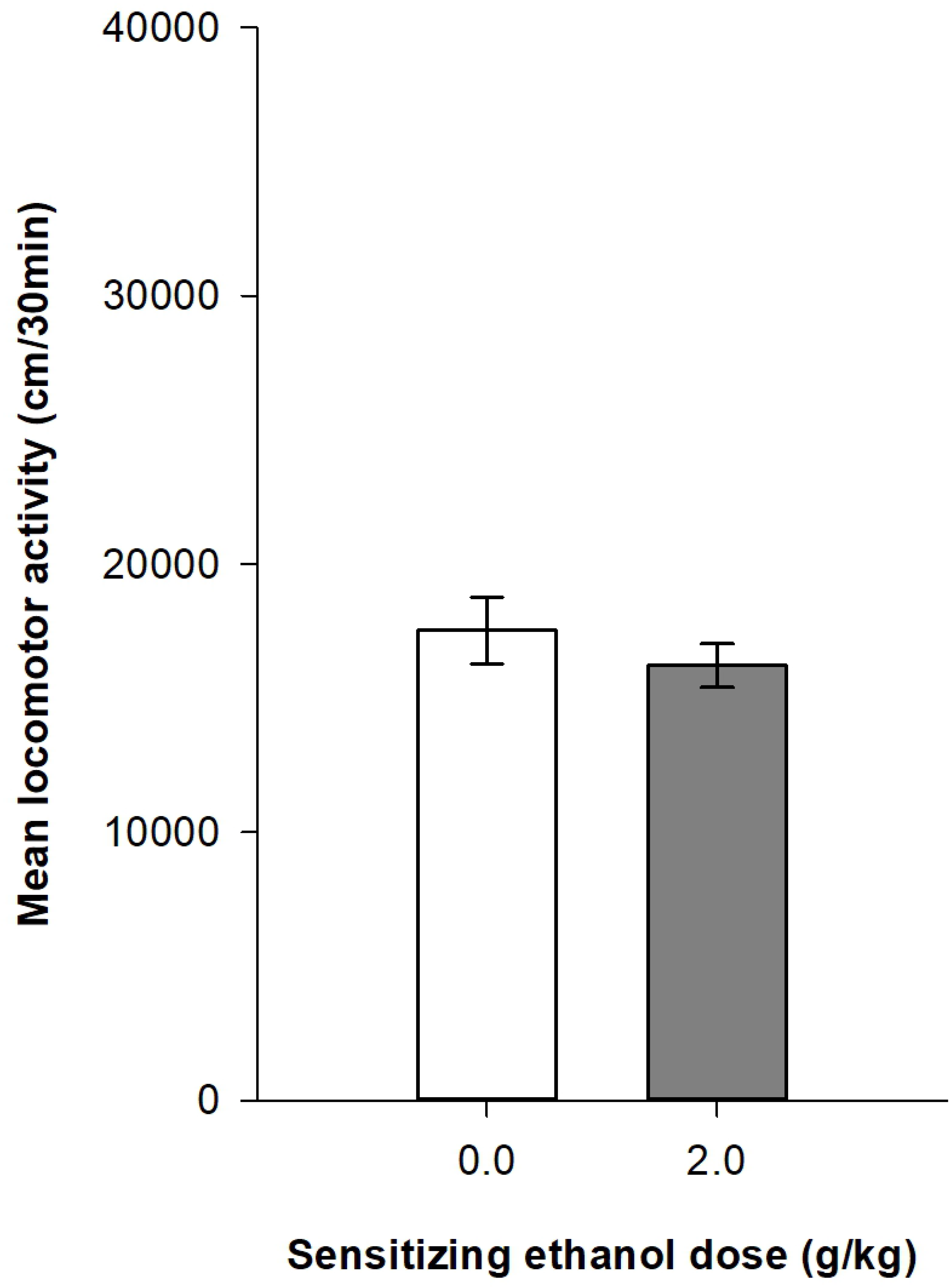
Total distance travelled during conditioned response test in DBA/2j Swiss mice. DBA/2j Swiss mice were administered with saline to test for the conditioned response induced by chronic ethanol administration (2.0 g/kg).

### Classifications of chronic ethanol-sensitized mice

#### Median split classification

Figures S1 and S2 (supporting information) show the development of ethanol sensitization respectively in Swiss and DBA2/J mice after their classification into respondent and resistant mice using the median split procedure described above. The mixed design ANOVAs show significant main effects for the group – control, resistant and respondent – (F_2,92_=156.96, p<0.0001, η²=0.66; F_2,62_=109.19, p<0.0001, η²=0.59 for Swiss and DBA2/J mice respectively) and for the session (45 sessions: F_44,4088_=52.14, p<0.0001, η²=0.19 ; F_44,2728_=11.57, p<0.0001, η²=0.099). There are also significant interactions group x session (F_88,4048_=21.94, p<0.0001, η²=0,17 ; F_88,2728_=3.36; p<0.0001, η²=0.06). Both groups of mice classified as resistant and respondent according to the median split procedure show significant increases in locomotor activity over ethanol sessions, although respondent mice reach significantly higher levels of ethanol-induced locomotor activity on the 45^th^ ethanol session as confirmed by the planned contrast (p<0.0001). However, it is noteworthy that even mice classified as “resistant to ethanol sensitization” developed significant levels of ethanol sensitization as confirmed by a significant difference between the 1^st^ and 45^th^ ethanol session in this group (within-subject sensitization). Interestingly, significant differences between mice classified as resistant and respondent are already observed on the first acute session (p<0.0001). In fact, resistant mice show a significantly weaker stimulant response to acute ethanol than respondent mice.

To test for the expression of ethanol sensitization, a one-way ANOVA is computed on the total locomotor activity during the test session (day 46). A significant effect of group is obtained for both Swiss (F_2,92_=95.92, p<0.0001, η²=0.68) and DBA2/J (F_2,62_=14.13, p<0.0001, η²=0.31) mice. Respondent mice show higher ethanol-induced locomotor activity than resistant mice.

However, resistant mice also show higher ethanol-induced locomotor activity than control mice, again indicating a significant ethanol sensitization in resistant mice.

As resistant and respondent mice significantly differ on the 1^st^ acute ethanol session, a rate of change between the first and last sessions was computed for each mouse as a ratio between locomotor activities on the 45^th^ and 1^st^ sessions. Such a ratio is usually used as an index of locomotor sensitization. One-way ANOVAs were computed on the rates of changes to test for significant differences between control, resistant and respondent mice. In Swiss mice, there is a significant effect of the group (F_2,92_=26.81, p<0.0001, η²=0.37), which is due to significant differences between mice treated with ethanol and the control group. In contrast resistant and respondent mice do not significantly differ as confirmed by post-hoc tests. In DBA/2J mice, there is no significant effect of the group on the rate of change in locomotor activity (F_2,62_=0.88, p=0.42, η²=0.028).

#### Extreme group classification

Figures S3 and S4 (supporting information) show the development of ethanol sensitization respectively in Swiss and DBA2/J mice after their classification into respondent and resistant mice using the extreme group procedure described above. The results obtained with this extreme group procedure are virtually identical to those obtained with the median split procedure. The mixed design ANOVAs show significant main effects for the group – control, resistant and respondent – (F_2,68_=222,893, p<0.0001, η²=0.77; F_2,44_=154.55, p<0.0001, η²=0.70 for Swiss and DBA2/J mice respectively) and for the session (45 sessions: F_44,2992_=46.21, p<0.0001, η²=0.24; F_44,1936_=8.37, p<0.0001,, η²=0.11). There are also significant interactions group x session (F_88,2992_=25,61, p<0.0001, η²=0.26; F_88,1936_=3.56; p<0.0001, η²=0.098). Similarly to the median split procedure, the planned contrasts indicate that respondent mice reach higher levels of ethanol-induced locomotor activity on the 45^th^ ethanol session, but that resistant mice develop significant levels of within-subject ethanol sensitization. Significant differences are also observed between mice classified as resistant or respondent on the 1^st^ acute ethanol session (p<0.0001), with respondent mice showing higher levels of ethanol-induced acute stimulant effects.

The one-way ANOVAs computed on the mean locomotor activity during the 30-min test session (day 46) show significant group effects in both Swiss (F_2,68_=116.47, p<0.0001, η²=0.77) and DBA2/J (F_2,44_=13.41, p<0.0001, η²=0.38) mice. Respondent mice show higher ethanol-induced locomotor activity than resistant mice, themselves showing significantly higher levels of activity than control mice.

Finally, the rate of change in locomotor activity was computed as the ratio between locomotor activities on the 45^th^ and 1^st^ sessions. In Swiss mice, there is a significant effect of the group (F_2,68_=25.30, p<0.0001, η²=0.43). As for the median split classification, this effect is explained by significant differences between ethanol and saline treated mice, whereas resistant and respondent mice do not differ in the rate of change. In DBA2/J mice, there is no significant effect of group in the rate of change (F_2,44_=1.24, p=0.299, η²=0.053).

#### D10/D1ratio classification

The D10/D1ratio classification leads to a very different pattern of results in comparison to the other two classifications. Figures S5A and S6A (supporting information) show the development of ethanol sensitization respectively in Swiss and DBA2/J mice after their classification into respondent and resistant mice using the procedure used by Botia and collaborators (10,19–21). The mixed design ANOVAs show significant main effects for the group – control, resistant and respondent – (F_2,92_=50.10, p<0.0001, η²=0.44 ; F_2,62_=33.80, p<0.0001, η²=0.40 for Swiss and DBA2/J mice respectively) and for the session (45 sessions: F_44,4048_=43.30, p<0.0001, η²=0.11 ; F_44,2728_=12.94, p<0.0001, η²=0.076). There are also significant interactions group x session (F_88,4048_=17.93, p<0.001, η²=0.094; F_88,2778_=3.72; p<0.0001, η²=0.045). Locomotor activity of both resistant and respondent mice from both strains gradually increase over repeated ethanol injections as confirmed by significant mean differences between the 1^st^ and 45^th^ ethanol sessions. Furthermore, the planned contrasts also reveal a significantly higher locomotor activity in respondent mice relative to resistant mice in both the 1^st^ acute session and the last 45^th^ ethanol session.

The one-way ANOVAs computed on the mean locomotor activity during the 30-min test session (day 46) show significant group effects in both Swiss (F_2,92_=69.95, p<0.0001, η²=0.60) and DBA2/J (F_2,62_=12.40, p<0.0001, η²=0.29) mice. However, this effect is due to a significant difference between the control group and both ethanol sensitized groups. In contrast to what is observed in the last 5-min ethanol session (session 45), the post-hoc tests on the 30-min test session fail to show a significant difference between resistant and respondent mice.

The one-way ANOVAs computed on the ratios between locomotor activities on the 45^th^ and 1^st^ sessions show significant main effects in both Swiss (F_2,92_=40,65; p<0.0001, η²=0.45) and DBA2/J (F_2,62_=20.13; p<0.0001, η²=0.39) mice. In both strains of mice, the post-hoc tests indicate significantly higher rates of change in respondent mice relative to resistant mice, while resistant mice significantly differ from control mice in Swiss mice, but not in DBA/2J mice.

Finally, a Cochran’s Q test was computed on the percentage of mice categorized as respondent when this classification procedure was used on the basis of locomotor activities at the 10^th^, 21^st^ and 45^th^ ethanol sessions. Whereas the percentage of respondent do not significantly change in DBA2/J mice (Q = 2.38, dl = 2, p < 0.30), there is a significant increase in the percentage of respondent mice in Swiss mice as the number of ethanol injections increase (Q = 21.7, dl = 2, p <0.0001). This indicates that some mice classified as resistant mice switch to the respondent group with additional ethanol injections.

## Discussion

### Development and expression of ethanol sensitization

To our knowledge, this was the first study showing the day by day development of ethanol sensitization following 45 ethanol injections in two strains of mice. Confirming previous reports, no tolerance to the stimulant effects of ethanol was observed even after a prolonged period of ethanol exposition (22,23). In contrast, the stimulant effects of ethanol continued to sensitize when the number of ethanol injections exceeded what was usually administered in previously published studies. As seen in Figures 1 and 4, ethanol sensitization can be described as a nearly logarithmic process divided in two phases. In a first phase of about 20 days, locomotor activity rapidly increased in both strains of mice over repeated ethanol injections. Then in a second phase, the development of ethanol sensitization decelerated in Swiss mice to reach a maximum level after about 40 ethanol injections, whereas ethanol sensitization seemed to have reached a stable ceiling effect in DBA2/J mice at that moment. At least in Swiss mice, the present results therefore showed that ethanol sensitization continue to progress significantly beyond the number of ethanol injections (8 to 20) used in most ethanol sensitization studies (4,10,30,40). However, it is noteworthy that the most important part of ethanol sensitization developed with the first 20 ethanol injections, such that previous studies reached the core phase of the development of ethanol sensitization. However, in previous studies, ethanol sensitization was often implicitly interpreted as a fully linear process. The results of the present study clearly show that it is not the case. In fact, the development of ethanol sensitization does not follow a linear development in time. Unfortunately, this temporal dimension of the development of ethanol sensitization is rarely discussed in the interpretation of ethanol sensitization results. The only exception is the differentiation of “acute sensitization” after a single ethanol injection from “chronic sensitization” after several ethanol injections (45–47). For example, mice are said to be non-sensitized or resistant to sensitization after a few ethanol injections. As shown in the present study, ethanol sensitization continues to develop over time in all mice, such that the term “delayed sensitization” might better characterize the so-called “non-sensitized mice”. It is currently unknown whether the speed of ethanol sensitization development is related to other components of alcohol abuse and addiction and should be the subject of further studies.

The 30-min test session also provided interesting observations about ethanol sensitization. In the control groups, naïve to ethanol before the test session, locomotor activity followed the typical time course effect after an acute ethanol administration. The acute locomotor effects of ethanol in mice were often described as biphasic (48,49). Rapidly after ethanol administration, locomotor activity significantly increased for a short period of time (usually 5-10 minutes) corresponding to the ascending limb of the blood alcohol concentration (50). Then locomotor activity returned to basal pre-injection levels or even occasionally to lower levels than the saline control group. This pattern of results was clearly apparent in Swiss mice (see control group in Figure 2A) with stimulant effects during the first 10-min after ethanol injection, whereas DBA2/J mice showed more prolonged stimulant effects for up to 30 minutes. Very interestingly, a long treatment with repeated intermittent ethanol injections led to a very significant ethanol sensitization characterized by both an upward shift of the magnitude of ethanol stimulant effects in both strains and a prolongation of these effects in time (up to 30 minutes) which was especially apparent in Swiss mice. On the sensitization test, ethanol sensitized mice from both strains expressed a strong ethanol-induced stimulation that was maintained throughout the test session. Another interesting observation came from the comparison of the two strains of mice. DBA2/J mice showed a higher stimulant response to ethanol than Swiss mice at the beginning of the sensitization procedure. This is probably the main reason why this strain of mice was so often chosen in ethanol sensitization studies. However, after a prolonged exposure to ethanol, Swiss mice finally reached similar levels of ethanol sensitization than DBA/2J mice on both the last ethanol sensitization session and the test session.

### Conditioned response induced by repeated ethanol injections

To test for the existence of an ethanol-induced conditioned response, all mice were injected with saline on the day following the sensitization test. In DBA2/J mice, no conditioned response was observed in spite of very strong levels of ethanol sensitizations that developed after 46 ethanol injections in the test context. In contrast, slightly increased levels of locomotor activity were observed in ethanol sensitized Swiss mice challenged with saline in the test context. However, this effect seems to be better interpreted as a lack of habituation to the test context rather than as an excitatory conditioned response. Indeed, a within-subject comparison between the habituation session (day 0) and conditioned test sessions (day 47) showed a significant decrease in locomotor activity in the saline control group, which is indicative of a habituation process, whereas locomotor activity remained stable in ethanol sensitized mice. Therefore, the significant difference between control and ethanol-sensitized Swiss mice on the conditioned test session was mainly due to a decrease of activity in the control group and not to an increase level of locomotor activity in ethanol sensitized mice. This supports the idea that ethanol prevented the process of habituation to the test environment.

A similar absence of ethanol-induced conditioned response was observed in other previously published studies, although it might have been attributable to an insufficient number of associations between the test context and ethanol injections. For example, an absence of a conditioned response was shown after 11 ethanol injections in DBA2/J mice (13,26). Similar results were also obtained in Swiss mice (27,51). In a previous study in our laboratory with a lower number of ethanol injections, no conditioned response was also observed in Swiss mice (4). Remarkably, this lack of higher locomotor activity in ethanol sensitized mice on the conditioned test was associated with a lack of significant habituation to the test context of the saline control group. These previous results further support the lack of habituation explanation. When the control mice do not show a significant habituation, probably due to an insufficient exposition to the test context, no differences are observed between ethanol sensitized and saline control mice on the conditioned test.

In contrast to the present results, some previous studies reported conditioned response in ethanol sensitized mice after a saline challenge (52–54). Surprisingly, these studies reported an excitatory conditioned response of very high magnitude, sometimes exceeding the sensitized response to ethanol (52) and sometimes higher than the levels of conditioned response observed with psychostimulants such as cocaine. The reasons for such discrepancies are difficult to explain and might be due to many methodological differences between studies. For example, Itzhak and Anderson used mice generated on a mixed C57BL⁄ 6J and SV129 background (53).

In conclusion, the present results extend previous reports indicating that sensitization to the stimulant effects of ethanol does not induce an excitatory conditioned response, even after 46 associations of a specific context with ethanol injections. The extensive exposition to ethanol in the present study makes it very unlikely that the lack of conditioned response was due to an insufficient pairing of the test context with ethanol injections.

### Classification of mice into resistant and respondent to ethanol sensitization

Despite recurrent criticisms from statisticians and methodologists (see for example (41–44)), it is common to categorize continous variables in various fields of science. In recent years such an approach was also used in ethanol sensitization studies (10,14,16,19–21,27–40). Three different procedures were used to classify mice after repeated ethanol injections into “ethanol sensitized” and “ethanol non sensitized” mice and into “resistant to ethanol sensitization” and “respondent to ethanol sensitization”. In the present study, these classification procedures were reexamined after a long ethanol sensitization procedure. A first striking conclusion of the present study is that the terms “ethanol non sensitized” and “resistant to ethanol sensitization” are inappropriate to qualify mice failing to show a significant sensitization after 10 to 20 ethanol injections. The present results clearly show that mice classified as “resistant to ethanol sensitization” according to these procedures developed very significant levels of ethanol sensitization when tested after 45 ethanol injections. In fact, these mice would be better characterized as showing a delayed development of ethanol sensitization.

When examining the patterns of results of mice after their classification, three dimensions in their behaviors were clearly apparent and must be considered in their characterization: the stimulant response to the first acute ethanol injection, the levels of sensitized ethanol stimulant effects at various ethanol sessions and the rate of change between the first and last ethanol injections. The first dimension characterizes the initial sensitivity of the mice to the stimulant effects of ethanol. The second dimension corresponds to the levels of stimulation experienced by the mice at a specific session and might be as important to explain individual mouse behavior at that time as the process of ethanol sensitization itself. Finally, the rate of change is the characteristic that best fits the definition of the ethanol sensitization process.

When comparing the three classifications, a very important point is that all of them led to groups of mice that already differ on the first dimension, i.e. the acute initial sensitivity to the stimulant effects of ethanol. This observation might have important consequences as such basal differences might well explain, at least in part, the differences in ethanol sensitization that will later occur. For example, a high rate of change (third dimension) is often more difficult to reach when the basal level of stimulation (first dimension) is already high. With the median split and extreme group classifications, respondent mice clearly show higher levels of acute ethanol stimulant effects than resistant mice. The respondent mice then maintain or even amplify this difference to keep higher levels at each time point throughout the sensitization procedure. However, the rate of change does not significantly differ between the two groups. One might therefore wonder whether such classifications do not simply identify mice that are differently sensitive to the stimulant effects of ethanol rather than to the sensitization process itself. It is noteworthy that the authors of such classifications report rates of sensitization that significantly differ between resistant and respondent mice in contrast to the present results (30,40). However, they did not use one of the measures that usually define ethanol sensitization, i.e. a score of difference between the last and the first sensitization sessions or a ratio between the last and first ethanol sessions. Instead they used a ratio between locomotor activities on the last ethanol sensitization session and the locomotor activity on the habituation session, i.e. a score of locomotor activity after a saline challenge. The use of such a ratio masks possible differences in the initial response to acute ethanol since this measure is not included in the ratio. This ratio directly derives from studies with psychostimulants. However, we believe that it is not appropriate to study ethanol sensitization as it does not take into account the fact that some mice show reduced levels of activity relative to their own basal levels of activity when injected with acute ethanol. Additionally, this ratio of the sensitized response divided by the basal locomotor activity without ethanol does not fit with the definition of sensitization, i.e. a progressive increase in the locomotor stimulant effects over repeated administrations of ethanol. This ratio does not characterize changes relative to the acute effects of ethanol but relative to a “normal” basal level of locomotor activity.

With the D10/D1 ratio classification, the reverse pattern of results is obtained on the acute ethanol session. Resistant mice show significantly higher ethanol-induced locomotor stimulant effects than respondent mice. Respondent mice then show a higher rate of increase to rise above the resistant mice after a few ethanol injections. However, the two groups of mice only maintain small differences in ethanol sensitization that tends to decrease with repeated ethanol injections. On the sensitization test (day 46), no statistically significant mean differences in the expression of ethanol sensitization can be detected between the groups of resistant and respondent mice. Additionally, when using this procedure of classification at various time points, the percentage of respondent mice significantly increase with the multiplication of ethanol injections from about 60% on the 10^th^ ethanol session to 90% on the last ethanol session. With additional ethanol injections, it might even be expected that the percentage of mice respondent to ethanol sensitization will eventually reach 100%. In a sense, the conclusion might be that with sufficient ethanol injections, all mice will eventually develop a sensitization to the effects of ethanol. This latter conclusion has the merit to show that the relative resistance/proneness to develop ethanol sensitization is dependent upon the number of ethanol injections and is not a stable trait. It is probably more appropriate to classify mice according to the speed at which they develop ethanol sensitization.

As a conclusion, the ideal classification of mice into resistant and respondent to ethanol sensitization should split the mice into two groups that would be characterized by an absence of differences in the acute response to ethanol (we want mice to differ on ethanol sensitization not on their initial response to the stimulant effects of ethanol), by differential rates of change (i.e. by differences in the development of ethanol sensitization) and therefore by the differential levels of sensitized ethanol stimulants effects that are achieved at some specific time points. Unfortunately, none of the reviewed classifications fulfills all these criteria. All three classifications involved significant differences in the initial acute stimulant effects of ethanol. The median split and extreme group classifications resulted in two groups of mice that differed in the levels of ethanol sensitized effects at many time points, including the sensitization test (day 46), but their rates of change were not significantly different. Finally, among those classifications, the D10/D1 ratio classification is probably the closest to reach all criteria. Although the groups differed on the rate of change and on the levels of ethanol sensitized effects at several time points, differences on this later criterion tended to decrease with the number of ethanol injections, such that no statistically significant differences were obtained on the test session after 45 ethanol injections. It is therefore very difficult to ascertain whether these classifications are specifically related to the sensitization process. They might all be more or less contaminated with other ethanol properties such as the initial sensitivity to ethanol stimulant or sedative effects. Finally, in agreement with previous warnings (41–44), we recommend avoiding the categorization of continuous variable and using the appropriate statistical tools for continuous variables. This is especially true with ethanol sensitization, which clearly shows several inter-related dimensions that must be included in the interpretation of the results.

## References

1. Kalivas PW, Stewart J. Dopamine transmission in the initiation and expression of drug- and stress-induced sensitization of motor activity. Brain Res Brain Res Rev. 1991 Dec;16(3):223–44.

2. Robinson TE, Berridge KC. The neural basis of drug craving: an incentive-sensitization theory of addiction. Brain Res Brain Res Rev. 1993 Dec;18(3):247–91.

3. Quoilin C, Didone V, Tirelli E, Quertemont E. Chronic ethanol exposure during adolescence alters the behavioral responsiveness to ethanol in adult mice. Behav Brain Res. 2012 Apr 1;229(1):1–9.

4. Didone V, Quoilin C, Tirelli E, Quertemont E. Parametric analysis of the development and expression of ethanol-induced behavioral sensitization in female Swiss mice: effects of dose, injection schedule, and test context. Psychopharmacology (Berl). 2008 Dec;201(2):249–60.

5. Hunt WA, Lands WE. A role for behavioral sensitization in uncontrolled ethanol intake. Alcohol Fayettev N. 1992 Aug;9(4):327–8.

6. Vanderschuren LJMJ, Pierce RC. Sensitization processes in drug addiction. Curr Top Behav Neurosci. 2010;3:179–95.

7. Newlin DB, Thomson JB. Chronic tolerance and sensitization to alcohol in sons of alcoholics: II. Replication and reanalysis. Exp Clin Psychopharmacol. 1999 Aug;7(3):234–43.

8. Newlin DB, Thomson JB. Alcohol challenge with sons of alcoholics: a critical review and analysis. Psychol Bull. 1990 Nov;108(3):383–402.

9. Didone V, Quoilin C, Dieupart J, Tirelli E, Quertemont E. Differential effects of context on psychomotor sensitization to ethanol and cocaine. Behav Pharmacol. 2016 Apr;27(2-3 Spec Issue):173–81.

10. Botia B, Legastelois R, Alaux-Cantin S, Naassila M. Expression of ethanol-induced behavioral sensitization is associated with alteration of chromatin remodeling in mice. PloS One. 2012;7(10):e47527.

11. Camarini R, Pautassi RM, Méndez M, Quadros IM, Souza-Formigoni ML, Boerngen-Lacerda R. Behavioral and neurochemical studies in distinct animal models of ethanol’s motivational effects. Curr Drug Abuse Rev. 2010 Dec;3(4):205–21.

12. de Araujo NP, Fukushiro DF, Grassl C, Hipólide DC, Souza-Formigoni MLO, Tufik S, et al. Ethanol-induced behavioral sensitization is associated with dopamine receptor changes in the mouse olfactory tubercle. Physiol Behav. 2009 Jan 8;96(1):12–7.

13. Legastelois R, Botia B, Naassila M. Blockade of ethanol-induced behavioral sensitization by sodium butyrate: descriptive analysis of gene regulations in the striatum. Alcohol Clin Exp Res. 2013 Jul;37(7):1143–53.

14. Quadros IMH, Hipólide DC, Frussa-Filho R, De Lucca EM, Nobrega JN, Souza-Formigoni MLO. Resistance to ethanol sensitization is associated with increased NMDA receptor binding in specific brain areas. Eur J Pharmacol. 2002 May 3;442(1–2):55–61.

15. Quadros IMH, Nobrega JN, Hipólide DC, de Lucca EM, Souza-Formigoni MLO. Differential propensity to ethanol sensitization is not associated with altered binding to D1 receptors or dopamine transporters in mouse brain. Addict Biol. 2002 Jul;7(3):291–9.

16. Quadros IMH, Nobrega JN, Hipolide DC, Souza-Formigoni MLO. Increased brain dopamine D4-like binding after chronic ethanol is not associated with behavioral sensitization in mice. Alcohol Fayettev N. 2005 Oct;37(2):99–104.

17. Quadros IMH, Souza-Formigoni MLO, Fornari RV, Nobrega JN, Oliveira MGM. Is behavioral sensitization to ethanol associated with contextual conditioning in mice? Behav Pharmacol. 2003 Mar;14(2):129–36.

18. Quoilin C, Didone V, Tirelli E, Quertemont E. Developmental differences in ethanol-induced sensitization using postweanling, adolescent, and adult Swiss mice. Psychopharmacology (Berl). 2012 Feb;219(4):1165–77.

19. Coune F, Silvestre de Ferron B, González-Marín MC, Antol J, Naassila M, Pierrefiche O. Resistance to ethanol sensitization is associated with a loss of synaptic plasticity in the hippocampus. Synap N Y N. 2017;71(2).

20. Legastelois R, Botia B, Coune F, Jeanblanc J, Naassila M. Deciphering the relationship between vulnerability to ethanol-induced behavioral sensitization and ethanol consumption in outbred mice. Addict Biol. 2014 Mar;19(2):210–24.

21. Botia B, Legastelois R, Houchi H, Naassila M. Basal anxiety negatively correlates with vulnerability to ethanol-induced behavioral sensitization in DBA/2J mice: modulation by diazepam. Alcohol Clin Exp Res. 2015 Jan;39(1):45–54.

22. Masur J, Boerngen R. The excitatory component of ethanol in mice: a chronic study. Pharmacol Biochem Behav. 1980 Dec;13(6):777–80.

23. Masur J, Oliveira de Souza ML, Zwicker AP. The excitatory effect of ethanol: absence in rats, no tolerance and increased sensitivity in mice. Pharmacol Biochem Behav. 1986 May;24(5):1225–8.

24. Brabant C, Quertemont E, Tirelli E. Influence of the dose and the number of drug-context pairings on the magnitude and the long-lasting retention of cocaine-induced conditioned place preference in C57BL/6J mice. Psychopharmacology (Berl). 2005 Jun;180(1):33–40.

25. Brabant C, Tambour S, Quertemont E, Ferrara A, Tirelli E. Do excitatory and inhibitory conditioning processes underlie psychomotor sensitization to amphetamine? An analysis using simple and multiple regressions. Behav Brain Res. 2011 Aug 1;221(1):227–36.

26. Meyer PJ, Phillips TJ. Bivalent effects of MK-801 on ethanol-induced sensitization do not parallel its effects on ethanol-induced tolerance. Behav Neurosci. 2003 Jun;117(3):641–9.

27. Abrahao KP, Quadros IMH, Souza-Formigoni MLO. Individual differences to repeated ethanol administration may predict locomotor response to other drugs, and vice versa. Behav Brain Res. 2009 Feb 11;197(2):404–10.

28. Abrahao KP, Ariwodola OJ, Butler TR, Rau AR, Skelly MJ, Carter E, et al. Locomotor sensitization to ethanol impairs NMDA receptor-dependent synaptic plasticity in the nucleus accumbens and increases ethanol self-administration. J Neurosci Off J Soc Neurosci. 2013 Mar 13;33(11):4834–42.

29. Abrahao KP, Quadros IMH, Andrade ALM, Souza-Formigoni MLO. Accumbal dopamine D2 receptor function is associated with individual variability in ethanol behavioral sensitization. Neuropharmacology. 2012 Feb;62(2):882–9.

30. Ferreira SE, Abrahao KP, Souza-Formigoni MLO. Expression of behavioral sensitization to ethanol is increased by energy drink administration. Pharmacol Biochem Behav. 2013 Sep;110:245–8.

31. Nona CN, Guirguis S, Nobrega JN. Susceptibility to ethanol sensitization is differentially associated with changes in pCREB, trkB and BDNF mRNA expression in the mouse brain. Behav Brain Res. 2013 Apr 1;242:25–33.

32. Nona CN, Li R, Nobrega JN. Altered NMDA receptor subunit gene expression in brains of mice showing high vs. low sensitization to ethanol. Behav Brain Res. 2014 Mar 1;260:58–66.

33. Nona CN, Bermejo MK, Ramsey AJ, Nobrega JN. Changes in dendritic spine density in the nucleus accumbens do not underlie ethanol sensitization. Synap N Y N. 2015 Dec;69(12):607–10.

34. Nona CN, Lam M, Nobrega JN. Localized brain differences in Arc expression between mice showing low vs. high propensity to ethanol sensitization. Pharmacol Biochem Behav. 2016 Mar;142:15–22.

35. Nona CN, Nobrega JN. A role for nucleus accumbens glutamate in the expression but not the induction of behavioural sensitization to ethanol. Behav Brain Res. 2018 15;336:269–81.

36. Abrahao KP, Souza-Formigoni MLO. Behavioral sensitization to ethanol results in cross-sensitization to MK-801 but not to NMDA administered intra-accumbens. Behav Brain Res. 2012 Dec 1;235(2):218–24.

37. Abrahao KP, Quadros IM, Souza-Formigoni MLO. Morphine attenuates the expression of sensitization to ethanol, but opioid antagonists do not. Neuroscience. 2008 Oct 28;156(4):857–64.

38. Abrahao KP, Goeldner FO, Souza-Formigoni MLO. Individual differences in ethanol locomotor sensitization are associated with dopamine D1 receptor intra-cellular signaling of DARPP-32 in the nucleus accumbens. PloS One. 2014;9(2):e98296.

39. Pildervasser JVN, Abrahao KP, Souza-Formigoni MLO. Distinct behavioral phenotypes in ethanol-induced place preference are associated with different extinction and reinstatement but not behavioral sensitization responses. Front Behav Neurosci. 2014;8:267.

40. Souza-Formigoni ML, De Lucca EM, Hipólide DC, Enns SC, Oliveira MG, Nobrega JN. Sensitization to ethanol’s stimulant effect is associated with region-specific increases in brain D2 receptor binding. Psychopharmacology (Berl). 1999 Oct;146(3):262–7.

41. MacCallum RC, Zhang S, Preacher KJ, Rucker DD. On the practice of dichotomization of quantitative variables. Psychol Methods. 2002 Mar;7(1):19–40.

42. Royston P, Altman DG, Sauerbrei W. Dichotomizing continuous predictors in multiple regression: a bad idea. Stat Med. 2006 Jan 15;25(1):127–41.

43. Altman DG, Royston P. The cost of dichotomising continuous variables. BMJ. 2006 May 6;332(7549):1080.

44. Irwin JR, McClelland GH. Negative Consequences of Dichotomizing Continuous Predictor Variables. J Mark Res. 2003 Aug 1;40(3):366–71.

45. Post RM, Weiss SR, Pert A, Fontana D. Conditioned components of cocaine sensitization. Clin Neuropharmacol. 1992;15 Suppl 1 Pt A:650A–651A.

46. Ceyhan M, Kayir H, Uzbay IT. Investigation of the effects of tianeptine and fluoxetine on pentylenetetrazole-induced seizures in rats. J Psychiatr Res. 2005 Mar;39(2):191–6.

47. Procópio-Souza R, Fukushiro DF, Trombin TF, Wuo-Silva R, Zanlorenci LHF, Lima AJO, et al. Effects of group exposure on single injection-induced behavioral sensitization to drugs of abuse in mice. Drug Alcohol Depend. 2011 Nov 1;118(2–3):349–59.

48. Durcan MJ, Lister RG. Time course of ethanol’s effects on locomotor activity, exploration and anxiety in mice. Psychopharmacology (Berl). 1988;96(1):67–72.

49. Smoothy R, Berry MS. Time course of the locomotor stimulant and depressant effects of a single low dose of ethanol in mice. Psychopharmacology (Berl). 1985;85(1):57–61.

50. Lewis MJ, June HL. Neurobehavioral studies of ethanol reward and activation. Alcohol Fayettev N. 1990 Jun;7(3):213–9.

51. Araujo NP, Fukushiro DF, Cunha JLS, Levin R, Chinen CC, Carvalho RC, et al. Drug-induced home cage conspecifics’ behavior can potentiate behavioral sensitization in mice. Pharmacol Biochem Behav. 2006 May;84(1):142–7.

52. Faria RR, Lima Rueda AV, Sayuri C, Soares SL, Malta MB, Carrara-Nascimento PF, et al. Environmental modulation of ethanol-induced locomotor activity: Correlation with neuronal activity in distinct brain regions of adolescent and adult Swiss mice. Brain Res. 2008 Nov 6;1239:127–40.

53. Itzhak Y, Anderson KL. Ethanol-induced behavioral sensitization in adolescent and adult mice: role of the nNOS gene. Alcohol Clin Exp Res. 2008 Oct;32(10):1839–48.

54. Itzhak Y, Martin JL. Blockade of alcohol-induced locomotor sensitization and conditioned place preference in DBA mice by 7-nitroindazole. Brain Res. 2000 Mar 10;858(2):402–7.

